# Cell type and locus-specific epigenetic editing of memory expression

**DOI:** 10.1101/2024.08.18.608454

**Authors:** Davide M. Coda, Lisa Watt, Liliane Glauser, Mykhailo Batiuk, Allison M. Burns, Cora L. Stahl, Johannes Gräff

**Affiliations:** Laboratory of Neuroepigenetics, Brain Mind Institute, School of Life Sciences, Ecole Polytechnique Federale Lausanne (EPFL); Lausanne, 1015, Switzerland; Bioinformatics Competence Center, Ecole Polytechnique Federale Lausanne (EPFL); Lausanne, 1015, Switzerland; Synapsy Research Center for Neuroscience and Mental Health Research, Ecole Polytechnique Federale Lausanne (EPFL); Lausanne, 1015, Switzerland

**Author notes:** University of Aberdeen; Aberdeen, AB24 3FX, Scotland, UK.

## Abstract

Epigenetic modifications parallel multiple memory processes, but evidence that the epigenetic makeup of a single site can guide learnt behaviors has so far been lacking. Here, we developed CRISPR-based epigenetic editing tools to address this question in a cell type-specific, locus-restricted and temporally controllable manner in the adult mouse brain. Focusing on learning-induced neuronal populations, we provide a proof-of-principle that site-specific epigenetic dynamics are causally implicated in memory expression.

## Main Text

Over the past decade, accumulating evidence has shown that memories are in part encoded in sparse populations of defined brain cells, so-called engrams^1^. Engram cells are nowadays the closest approximation of a memory trace, i.e., the physical substrate of a memory: Activated by learning, they display enhanced structural and synaptic plasticity^2^, regulate memory retrieval^3^, and are of lasting nature^4^.

At the same time, multiple studies have revealed that epigenetic mechanisms – mainly histone acetylation and DNA methylation – may contribute to memory formation, storage and change^5^. However, experimental evidence in support of these findings has been obtained by wide-range pharmacological agents, by genetic approaches that modify enzymes responsible for epigenetic changes, and by their study in either whole-tissue homogenates or broadly defined cell types such as excitatory neurons^6^. It is only recently that epigenetic modifications were found to occur in engram cells following learning^7^, but a causal, locus-restricted interrogation thereof for regulating memory expression remains to be demonstrated.

To address this question, we developed CRISPR-dCas9-based epigenetic editing tools^8–10^ for cFos-driven engram tagging technologies^11^ that allow for the induction of locus-specific chromatin alterations within sparse memory-bearing neuronal ensembles. As editing target, we focused on *Arc*, an immediate early gene (IEG) known for its role in learning and synaptic plasticity^12,13^, whose promoter region shows primed, i.e., increased chromatin accessibility compared to other IEGs in the mouse dentate gyrus (DG) (Extended Data Fig. 1). Since primed chromatin is a more dynamic epigenetic state than closed chromatin^14^, the *Arc* promoter represents an ideal site to probe for the importance of locus-specific epigenetic regulation.

To assess the behavioral consequences of epigenetically editing the *Arc* promoter within engram cells, we first engineered the epigenetic repressor dCas9-KRAB-MeCP2^9^ in a spatiotemporally regulatable manner, namely by combining it with an OFF doxycycline (DOX)-controllable tetracycline responsive element (TRE) in a lentiviral construct (Fig. 1A). We stereotaxically delivered this construct into the DG of cFos-tTA mice, which express the tetracycline-controlled transactivator (tTA) upon learning^11^. We used a second lentivirus expressing five U6-driven single-guide RNAs (sgRNAs) targeting the *Arc* promoter, and as controls we expressed U6-driven non-targeting (NT) sgRNAs.

**Fig. 1.**
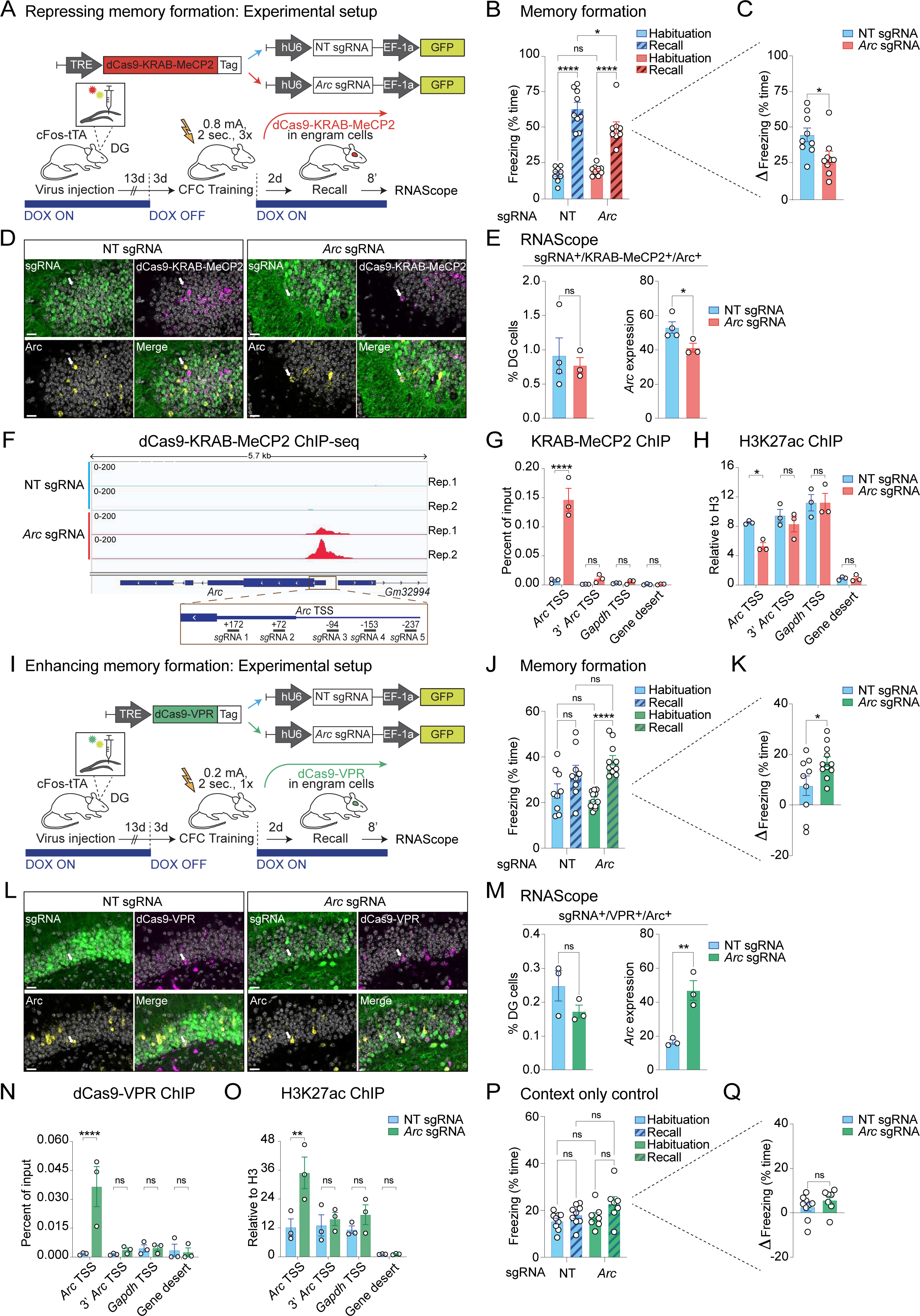
Bidirectional epigenetic editing of memory formation. **(A)** Schematic (top) and experimental design (bottom) for dCas9-KRAB-MeCP2-based epigenetic editing of the *Arc* promoter in cFos-tTA mice. For abbreviations, see text. (**B-C**) Compared to animals injected with NT sgRNA (n = 9), dCas9-KRAB-MeCP2 animals injected with *Arc* sgRNAs (n= 9) showed impaired memory formation. Data are means ± SEM compared either by (B) one-way ANOVA with Tukey9s post hoc test, or by (C) a one-tailed, unpaired t test. (**D**) Representative confocal images showing mRNA expression levels of *Arc* (yellow) and dCas9-KRAB-MeCP2 (magenta) alongside with NT or *Arc* sgRNA (GFP, green) in the DG. Scale bars, 20 μm. Arrows indicate neurons positive for dCas9-KRAB-MeCP2, sgRNA and *Arc*. (**E**) Quantification of the number of DG cells triple positive for dCas9-KRAB-MeCP2^+^/sgRNA^+^/*Arc+* and the level of *Arc* expression in the *Arc* sgRNA (n = 4) and NT sgRNA group (n = 3). Data are means ± SEM compared by a two-tailed, unpaired t test. (**F**) IGV genome browser of the *Arc* locus displaying ChIP-seq tracks for dCas9-KRAB-MeCP2 from N2A cells co-transfected with *Arc* or NT sgRNA in two independent biological replicates (Rep). Bottom panel, enlarged view of the region surrounding the *Arc* promoter with the genomic sequences targeted by the sgRNAs highlighted. (**G-H**) ChIP-qPCR on N2A cells treated as in F showing occupancy of (G) dCas9-KRAB-MeCP2 and (H) H3K27ac at the transcription start site (TSS) of *Arc*. Plotted are means ± SEM of three independent experiments compared by two-way ANOVA with Sidak9s multiple comparisons test. (**I**) Schematic (top) and experimental design (bottom) for dCas9-VPR-epigenetic editing of the *Arc* promoter in cFos-tTA mice. (**J-K**) Compared to animals injected with NT sgRNA (n = 9), dCas9-VPR plus *Arc* sgRNA injected animals (n = 11) showed improved memory recall. Data are means ± SEM compared either by (J) one-way ANOVA with Tukey9s post hoc test or by (K) a one-tailed, unpaired t test. (**L**) As in D, except that dCas9-VPR is expressed instead. Scale bars, 20 μm. Arrows indicate neurons positive for dCas9-VPR, sgRNA and *Arc*. **(M**) Quantification of the number of DG cells triple positive for dCas9-VPR^+^/sgRNA^+^/*Arc+* and the level of *Arc* expression in the *Arc* sgRNA (n = 3) and NT sgRNA group (n = 3). Data are means ± SEM compared by a two-tailed, unpaired t test. **(N-O)** As in G-H, except that dCas9-VPR is expressed instead. For N and O, plotted are means ± SEM of three independent experiments compared by two-way ANOVA with Sidak9s multiple comparisons test. (**P-Q**) As in I, except that no shock was administered during context exposure on the training day for mice injected with dCas9-VPR plus NT sgRNA (n = 9) or the Arc sgRNA (n = 7). Data are means ± SEM compared either by one-way ANOVA with Tukey9s post hoc test (P) or by a one-tailed, unpaired t test (Q). For all figure panels: ns, not significant; *P < 0.05, **P < 0.01 and ****P < 0.0001.

Then, to trigger dCas9-KRAB-MeCP2 expression in engram cells activated by learning, cFos-tTA mice were taken off DOX 3 days prior to contextual fear conditioning (CFC), an associative memory task in which mice learn to pair a specific context with an unpleasant experience (i.e., an electrical footshock). Immediately after CFC, mice were put on DOX again to prevent learning-unrelated TRE expression. 2 days later, we measured the animals9 memory performance by re-exposing them to the conditioned context in the absence of footshock (Fig. 1A). We found that dCas9-KRAB-MeCP2 plus *Arc* sgRNA mice exhibited significantly less freezing (i.e., the learnt association between footshock and context) than mice expressing NT sgRNA (Fig. 1B), indicating reduced memory formation. Importantly, normalizing the freezing rates during the recall phase to those during pre-conditioning (& freezing) confirmed that the reduction in memory expression was not due to alterations in baseline freezing behavior (Fig. 1C). Moreover, measurements of locomotion (distance travelled) and anxiety (time spent in the inner region of the behavioral apparatus) did not differ between the two groups (Extended Data Fig. 2, A and B), excluding memory-unrelated effects of the dCas9-KRAB-MeCP2 manipulation. At the molecular level, *Arc* expression in dCas9-KRAB-MeCP2-expressing cells was reduced in the *Arc* sgRNA compared to the NT group, while the percentage of DG cells expressing both the dCas9-KRAB-MeCP2 and the sgRNA vector was similar between the two groups (Fig. 1, D and E, and Extended Data Fig. 2C). Epigenetically, dCas9-KRAB-MeCP2 binding to the *Arc* promoter decreased the occupancy of the active chromatin mark H3K27ac, and the downregulation of *Arc* mRNA was paralleled by few off-target effects (Fig.1, F-H, and Extended Data Fig. 3, A-D) as assessed in N2a cells.

Next, we tested whether an epigenetic activation of *Arc* would lead to the opposite behavioral effect. To this end, we stereotaxically injected an OFF DOX-inducible version of the epigenetic activator dCas9-VPR^8,10^ (i.e., TRE-dCas9-VPR), as well as lentiviruses containing *Arc* or NT sgRNAs into the DG of cFos-tTA mice (Fig. 1I). We used the same experimental timeline as before, except for employing a subthreshold CFC protocol^15^, which did not lead to memory formation in the NT control group (Fig. 1J). In contrast, we found that dCas9-VPR plus *Arc* sgRNA mice showed a significant increase in freezing at recall, indicating improved memory formation (Fig. 1, J and K). No differences in locomotion nor baseline behavior were observed between the two groups (Extended Data Fig. 2, D and E). Molecular analyses revealed similar infection efficiency, but higher levels of *Arc* in engram cells positive for dCas9-VPR and *Arc* sgRNA compared to NT controls (Fig. 1, L and M, and Extended Data Fig. 2F), while epigenetically, dCas9-VPR binding to the *Arc* promoter increased H3K27ac and *Arc* mRNA with no off-target effects (Fig. 1, N and O, and Extended Data Fig. 3, E-G). Furthermore, context exposure without conditioning did not lead to increased freezing in dCas9-VPR plus *Arc* sgRNA mice, testifying to the specificity of the dCas9-VPR-based memory enhancement to the fearful experience (Fig. 1, P and Q, and Extended Data Fig. 4, A and B). Together, these experiments suggest that the epigenetic makeup of a single locus within sparse DG engram cells is necessary and sufficient to regulate memory expression.

In a second step, we asked whether the epigenetic editing effects on behavior are reversible within subject, which serves the purpose to examine their plasticity, a defining feature of any memory-related process. To this end, we employed the anti-CRISPR protein AcrIIA4^16^, which functions by occluding dCas99s PAM recognition domain, hence negating dCas9 binding to the DNA (Fig. 2A), and rendered it inducible by coupling it to a DOX ON controllable TRE promoter (Fig. 2B). *In vitro*, we found that AcrIIA4 induction reverted the dCas9-VPR-mediated increase of *Arc* within 3 days (Fig. 2, C and D, and Extended Data Fig. 5, A and B).

**Fig. 2.**
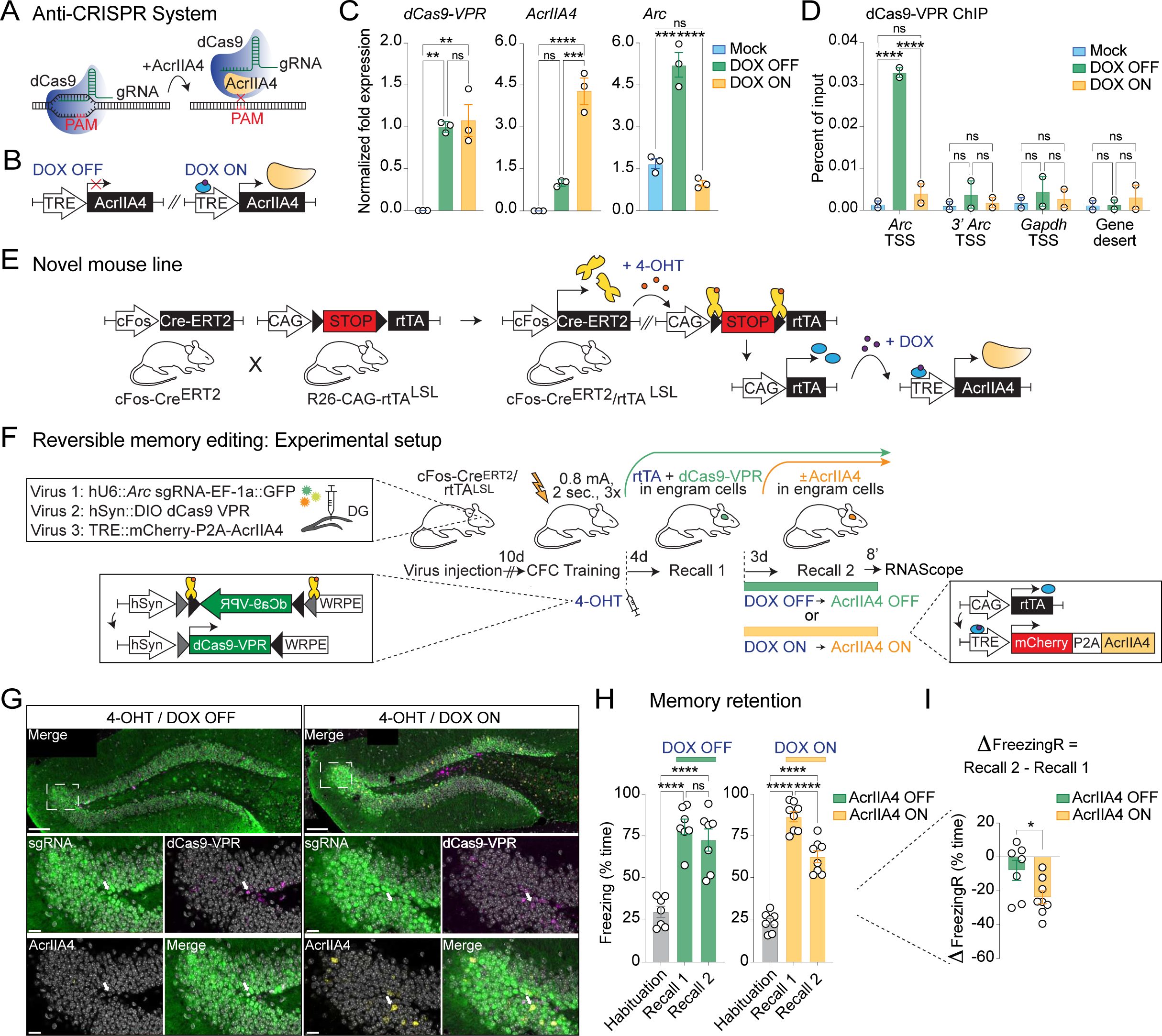
Within subject reversible epigenetic memory editing. (**A-B**) Schematic of the mechanism of action of the anti-CRISPR protein AcrIIA4 (A), which was rendered inducible by combining it with DOX ON activated TRE promoter (B). (**C**) qPCR on N2A cells transfected with dCas9-VPR, *Arc* sgRNA and AcrIIA4 showing that AcrIIA4 induction abolishes the dCas9-VPR mediated increase in *Arc* mRNA. Data are means ± SEM of three independent experiments compared by one-way ANOVA with Tukey9s post hoc test. (**D**) ChIP-qPCR on N2A cells treated as in C showing occupancy of dCas9-VPR at the transcription start site (TSS) of *Arc*. Plotted are means ± SEM of two independent experiments compared by two-way ANOVA with Tukey9s multiple comparisons test. (**E**) Breeding schematic of the novel double transgenic mouse line cFos-Cre^ERT2^/rtTA^LSL^. (**F**) Experimental timeline using cFos-Cre^ERT2^/rtTA^LSL^ the line to uncouple the expression of dCas9-VPR (4-OHT dependent) from the expression of AcrIIA4 (DOX ON-dependent). (**G**) Representative confocal images showing the expression of dCas9-VPR mRNA (magenta) alongside with *Arc* sgRNA (GFP, green) and the mCherry mRNA reporter for AcrIIA4 (yellow) in the DG. Scale bars, 100 μm. Smaller panels show enlarged areas within the dotted squares. Scale bars, 20 μm. Arrows indicate neurons positive for either dCas9-VPR and *Arc* sgRNA (4-OHT / DOX OFF) or dCas9-VPR, *Arc* sgRNA and AcrIIA4 (4-OHT / DOX ON). (**H-I**) Compared to animals expressing dCas9-VPR plus *Arc* sgRNA and kept off DOX (n = 7), animals put ON DOX after the first recall to induce AcrIIA4 (n = 8) showed impaired memory retention, expressed as freezing percentage (H) and the freezing differential between the second and first recall (I). Data are means ± SEM compared either by (H) one-way ANOVA with Tukey9s post hoc test or by (I) a one-tailed, unpaired t test. For all figure panels: ns, not significant; *P < 0.05, **P < 0.01, ***P < 0.001 and ****P < 0.0001.

To use this setup *in vivo,* we designed a novel double transgenic labeling system that allows for the sequential induction of dCas9-VPR and AcrII4 in the same engram cells. Specifically, we crossed cFos-Cre^ERT2^ mice^4^, in which tamoxifen-dependent Cre-recombinase expression is restricted to engram cells, with R26-CAG-rtTA^LSL^ mice^17^, which contain a loxP-flanked reverse tetracycline controlled transactivator (rtTA) that binds the TRE promoter in a DOX-ON dependent manner (Fig. 2E). In the resulting double transgenic cFos-Cre^ERT2^/R26-CAG-rtTA^LSL^ mice (hereof referred to as cFos-Cre^ERT2^/rtTA^LSL^), we stereotaxically delivered a Cre-dependent version of dCas9-VPR (using a double inverted open reading frame (DIO)-dCas9-VPR), AcrIIA4 under the control of TRE, and the *Arc* sgRNAs to their DG (Fig. 2F). Immediately after CFC, cFos-Cre^ERT2^/rtTA^LSL^ mice received an injection of 4-OHT, which triggered the expression of both rtTA and dCas9-VPR in learning-activated neurons. 4 days later, memory formation was measured with a first recall session as before. Thereafter, one group of mice received DOX, which induced AcrIIA4 expression, while the other group of mice did not receive DOX, thus AcrIIA4 was not expressed (Fig. 2G). Another 3 days later, all mice underwent a second recall test, by which their memory retention could be assessed.

For mice in which dCas9-VPR but not AcrIIA4 was induced (i.e., the DOX OFF group), we found that freezing levels on the second recall did not significantly differ from those of the first recall (Fig. 2H). Since mice without dCas9-VPR induction show reduced freezing upon the second memory recall (Extended Data Fig. 6), this finding indicates that dCas9-VPR induction leads to improved memory retention, in line with dCas9-VPR facilitating memory formation (Fig. 1K). Conversely, for animals in which dCas9-VPR induction was followed by AcrIIA4 activation (i.e., the DOX ON group), we observed reduced freezing at the second recall (Fig. 2H, I), indicating that the improved memory retention upon dCas9-VPR was negated. Importantly, AcrIIA4 induction did not alter basal locomotion nor anxiety levels (Extended Data Fig. 5, C and D). These reversibility experiments show that the observed behavioral effects are a direct consequence of the epigenetic regulation of *Arc*, which is thus plastic within the same animal.

In summary, by engineering a spatiotemporally inducible CRISPR-based epigenetic editing system in engram cells, we found that the epigenetic makeup of a single locus in a sparse neuronal population can bidirectionally regulate memory expression in reversible manner. From a neurobiological perspective, these experiments provide the hitherto finest-grained evidence of how epigenetic dynamics can impact memory capacities. With the caveat that we mechanistically interrogated only one site in the genome, in one brain region, in one memory task and only in male animals, these findings indicate that chromatin alterations might act as functionally relevant mnemonics to regulate the expression of learnt information.

From a methodological perspective, this study further illustrates how epigenetic editing tools, including the anti-CRISPR protein AcrIIA4, can be directed to designated cell populations in the adult mouse brain. Given its modular set-up, similar experimental approaches may in future studies prove useful to better understand other phenotypes from the perspective of cell and locus-specific epigenetic regulation: Within the field of memory research, for disorders characterized by aberrant memory processing; beyond the field of memory research, where epigenetic mnemonics have been proposed for drug-related memories^18^ as well as for stress-associated behaviors following childhood trauma^19^; and also outside the field of neuroscience, where cellular memories established during development^20^, following inflammation^21^ or after immune activation^22^ are known to be epigenetically encoded.

## Methods

### sgRNA design and cloning

To design guide sequences for the CRISPR-dCas9 systems a total of 439 bp spanning the promoter of the mouse gene *Arc* were used as input for the CRISPROR program (http://crispor.tefor.net)^23^. Five sgRNAs were selected and DNA oligonucleotides corresponding to the chosen sgRNA sequences (see Table S2) were individually cloned into pLenti SpBsmBI sgRNA Hygro (Addgene 62205) containing an U6 driven sgRNA scaffold. Afterwards, the individual sgRNAs were multiplexed in the same plasmid using NEBridge Golden Gate Assembly kit (NEB) with custom designed primers and cloned into the lentiviral pVLX-EF1a-GFP backbone. The sgRNA sequence against the bacterial *LacZ* gene was used as a non-targetting (NT) control sgRNA as previously described^24^. The plasmids for dCas9-VPR and dCas9-KRAB-MeCP2 were generated in-house by subcloning the lenti-EF1a-dCas9-VPR-Puro (Addgene 99373) and the lenti-SYN-dCas9-KRAB-MeCP2 (gift from Dr. Jeremy Day) into the lentiviral pVLX backbone under control of either the CMV promoter (*in vitro* experiments) or the TRE (*in vivo* experiments). For the AcrIIA4-mediated reversibility experiment: The pVLX-Dio-hSyn-dCas9-VPR plasmid was generated in-house by subcloning the dCas9-VPR sequence into the pVLX-Dio-hSyn backbone; the pVLX-hU6-ArcsgRNAs-EF1a-GFP-TRE-mCherry-P2A-AcrII4A (*in vitro* experiments) and the pVLX-TRE-mCherry-P2A-AcrII4A (*in vivo* experiments) were generated starting from pHR-Ef1a-mCherry-P2A-AcrIIA4 (Addgene 125148); the pSIN-hPGK-rtTA plasmid was a gift from Dr. Didier Trono. A full list of the plasmids used in the study is provided in Table S2.

### Lentivirus production

Lentiviral vectors were produced as previously described^25^. Briefly, HEK-293T cells (ATCC) were transfected using calcium phosphate with a second-generation packaging system composed of the pMD2G and psPAX2 vectors alongside the different pVLX vectors used in this study. After 4 days, medium was collected and centrifuged in an ultracentrifuge at 19,000 r.p.m. for 90 minutes at 4 °C. The pellet was resuspended in 1× PBS pH 7.4 and 0.5% BSA and viral titer was determined using the HIV-1 p24 antigen ELISA kit (Zeptometrix Corp). Injections were performed with 500 ng of each virus, which corresponded to a volume between 200 and 500 nL depending on the individual virus titer.

### Cell culture and transfection

N2A cells were obtained from ATCC, banked at EPFL and cultured in Dulbecco9s Modified Eagle Medium GlutaMAX# (Gibco) supplemented with a 1% Penicillin-Streptomycin solution (Gibco) and 10% Fetal Bovine Serum (Gibco). Cells were transfected with the appropriate plasmids using Lipofectamine 2000 (Invitrogen) according to manufacturer9s instructions for 48 h before being processed for downstream analyses. To induce the expression of the AcrIIA4 construct, cells were treated with 1)ug/ml of doxycycline.

### Immunocytochemistry

N2A cells were cultured on coverslips previously coated with polyD-Lysin (Corning) and transfected as described above. After 48 h of incubation, cells were fixed for 12 minutes at room temperature (RT) in a 4% paraformaldehyde, 4% sucrose PBS-based solution. Next, samples were incubated in a humidity chamber overnight at 4 °C with the primary antibody. The following day, cells were washed with 1x PBS and incubated with the secondary antibody for 1 h at RT. Finally, coverslips were mounted on a slide with Vectashield-DAPI and images acquired using an Olympus Slide Scanner (Olympus VS120) at 10X magnification. Antibodies used are listed in Table S2.

### RNA extraction and q-PCR

Total RNA was extracted using Trizol (Thermo Fisher Scientific), and cDNA synthesized using the SuperScript III First-Strand Synthesis kit (Invitrogen) according to manufacturer9s instructions. All qPCRs were performed with the FAST SYBR Green Master Mix (Applied Biosystems) with 300 nM of each primer and 2 µl of diluted cDNA. Each biological sample was loaded in technical triplicates and fluorescence acquisition was carried out on a StepOnePlus Real-Time PCR System (Applied Biosystems). Calculations were performed using the ——Ct method and levels of *Gapdh* mRNA and *Actin* mRNA as reference genes. Unless otherwise stated in the figure legends, means ± SEM from at least three independent biological experiments are shown. Primer sequences are listed in Table S2.

### Chromatin immunoprecipitation (ChIP)

Samples for ChIP-qPCR were obtained as described previously with minor adaptations^26^. Briefly, 20 million cells were crosslinked using 1% formaldehyde for 10 minutes at RT, followed by lysis in 5 mM HEPES (pH 8), 85 mM KCl, 0.5% NP-40. After centrifugation, nuclei were washed in 5 mM HEPES (pH 8), 85 mM KCl and then sonicated to 100–400 bp using 15 cycles 30= on-off in 1ml of 50 mM Tris-HCl (pH 8), 10 mM EDTA, 1% SDS using a Covaris LE220 sonicator. The resulting chromatin was diluted 5 times, pre-cleared with protein A or G Dynabeads (Invitrogen) and then incubated overnight with antibodies in immunoprecipitation (IP) buffer (50 mM HEPES (pH 7.5), 1% Triton X-100, 150 mM NaCl). The amount of sonicated chromatin used for each IP corresponded to 2 million cells for the H3K27ac or total H3 IP and to 10 million for the dCas9 IP. Protein A or G Dynabeads (100 µl) blocked with BSA were then added for 6 h, followed by six washes in IP buffer containing 0.1% sodium deoxycholate, 0.1% SDS and 1 mM EDTA, with the last three additionally containing 500 mM NaCl. A wash was then performed with 10 mM Tris-HCl pH 8, 250 mM LiCl, 1 mM EDTA, 1% NP-40 and 0.5% sodium deoxycholate followed by a final wash in 10 mM Tris-HCl pH 8, 1 mM EDTA. Chromatin was eluted in 1% SDS, 0.1 M NaHCO3. De-crosslinking and Proteinase K digestion was carried out at 65°C overnight. The resulting enriched chromatin was cleaned up using a QiaQuick PCR purification kit (Qiagen) according to manufacturer9s instructions. qPCR was carried out as for gene expression analysis, with a standard curve consisting of 6 points and three-fold dilutions. Values for each IP sample were normalized relative to corresponding input chromatin for the same treatment. Primer sequences and antibodies are listed in Table S2.

### ChIP-seq library preparation and analysis

Samples for ChIP-seq with the corresponding inputs were prepared as described above for ChIP-qPCR. Next, NEBNext Ultra II DNA Library Preparation Kit (NEB) was used to generate the libraries according to manufacturer9s instructions. Samples were then multiplexed and 75 bp pair end reads were generated on a Nextseq500 (Illumina). After reads were trimmed for NEBNext Ultra II DNA (TruSeq) adaptors, the FastQ files were demultiplexed using bclconvert (v3.9.3, Illumina) and aligned to the mm10 genome using bowtie2 (v2.4.5) in paired end mode and using default parameters. The R library Csaw (v.1.30.1) was used for peak calling, with peaks being considered non-significant for regions that had less than a three-fold enrichment compared to their 2 kb neighbourhood, and reads were normalized using the locally estimated scatterplot smoothing (loess) algorithm. Then, peaks were assigned to the nearest promoter with the distanceToNearest function from GenomicRanges. Finally, peaks in the immunoprecipitated samples were compared to their respective input samples by performing a differential enrichment analysis using EdgeR (v3.38.4). The threshold for significance was set at logFC > 2 and p-value <0.05. When assigning peaks to genomic features promoter regions were defined as 2kb upstream and 200bp downstream of the gene TSS. ChIP-seq data were visualized using the Integrative Genomics Viewer (IGV, version 2.16.0).

### Animals

All animals and procedures used in this study were approved by the Veterinary Office of the Federal Council of Switzerland under the animal experimentation license VD2808.2. cFos-tTA male mice were bred in house from the original JAX strain #018306 on a C57Bl/6JR background. Animals were group-housed in a 12 h light/dark cycle with water and food available ad libitum. DOX (0.2 mg/ml) was given orally through the water supply beginning at least 7 days before the start of the experiment, and only ceased 3 days prior to the session where the tagging window was desired. DOX was then provided back to the animals as soon as the tagging window was no longer needed. Double transgenic cFos-Cre^ERT2^/R26-CAG-rtTA^LSL^ animals were generated in house by crossing the original JAX strains #030323 and #029617. cFos-Cre^ERT2^/R26-CAG-rtTA^LSL^ male mice were injected i.p. with tamoxifen (4-hydroxytamoxifen, Sigma-Aldrich, 50 mg/kg) immediately after CFC. Tamoxifen was prepared as follows: Powdered tamoxifen was dissolved in ethanol 100% at a concentration of 20 mg/mL and stored at –20°C. On the day of the experiment, tamoxifen was re-dissolved by shaking at 37°C, then 2 volumes of corn oil were added and ethanol was evaporated shaking at 37°C for a final concentration at 10 mg/mL. Tamoxifen was kept at 37°C until injection to prevent precipitation. Mice were between 8-13 weeks old at the start of the experiments, and all males. All behavioral procedures were performed between 1 pm and 5 pm local time and animals were randomly assigned to experimental groups.

### Behavioural procedures

Contextual fear conditioning behavioural experiments were performed using a TSE Multi Conditioning System. CFC encoding consisted in a first 3 minute exploration phase, followed by 2 s long foot shocks spaced by 28 s. The number of repetitions of the pause-shock stage as well as the electrical current varied depending on the experiment. Briefly, the subthreshold CFC paradigm consisted in one (Fig. 1I) 2 s long shocks of 0.2 mA and the strong paradigm of three 2 s long shocks of 0.8 mA (Fig. 1A, Fig. 2E and Extended Data Fig. 6). No shock control animals underwent the same procedure but did not receive any shocks (Fig. 1, P and Q). After the last shock, the animal was left in the chamber for an additional 15 s and brought back to its home cage. The recall phase consisted in a 3 minute exposure to the same context, without any shock. The time after encoding at which recall took place varied depending on the experiment. The movement of the animals was automatically measured using an infrared beam cut detection system (TSE Systems) and freezing was calculated as the absence of movement for more than 500 ms.

### Viral injections

Surgeries were performed as described previously with minor adaptations^27^. Before each surgical procedure, mice were injected intraperitoneally (IP) with an anaesthetic mix of fentanyl (0.05 mg/kg), midazolam (5 mg/kg), and metedomidin (0.5 mg/kg), followed by a subcutaneous injection of an analgesic mix (lidocaine 6 mg/kg and bupivacaine 2.5 mg/kg) at the surgery site. Animals were then placed on a stereotaxic micromanipulator frame (Kopf Instruments), skin disinfected with betadine and opened with a scalpel. To target the DG, holes were drilled in the skull with a 30-gauge drill bit at ±1.3 mm medio-lateral, -2.0 mm anterior-posterior. A virus-loaded micropipette (BLAUBRAND, intraMARK, tip diameter 10-20 µm) was lowered to -2.0 mm and 500 ng of viral particles were injected bilaterally at a flow rate of 0.2 µl/minute. After 5 minutes, the micropipette was raised 20 µm from the target for a further minute to prevent backflow and allow diffusion, then removed from the brain at 10 µm/s. After the injections, the skin was sutured and atipamezol (2.5 mg/kg) was administered IP to reverse the anesthesia. Post-surgery, mice were moved to a clean cage on a heated pad until they regained mobility, and ultimately returned to their home cage with paracetamol provided in the drinking water for a week (500 mg per 250 ml). After sacrificing the mice and processing the brains as described below, surgery efficiency was verified for each animal by visualizing the GFP signal and mis-injected brains were excluded from behavioural and histological analyses.

### Histology and RNA scope

8 minutes after the last behavioral test, animals were anesthetized with pentobarbital (150 mg/kg) and transcardially perfused with first PBS and then 4% paraformaldehyde (PFA) in PBS. Brains were extracted, post-fixed overnight in 4% PFA, transferred in a cryoprotectant solution (30% sucrose in PBS) for at least 48 h, and frozen at -80°C. Sections of 20 μm were cut using a cryostat, and kept at -20°C in an antifreeze solution (30% ethylene glycol, 15% sucrose, 0.02% azide in 1× PBS) until staining. RNA scope followed by immunohistochemistry was performed using the RNA Scope Multiplex Fluorescent V2 kit (ACD Bio) according to manufacturer9s instructions. Briefly, mounted slices were incubated 30 minutes at 60°C, fixed for 15 minutes at 4°C in 4% PFA and dehydrated by incubation in increasing ethanol concentrations (50%, 70%, 100%). Then, a quenching step was performed (H_2_O_2_ for 10 minutes at RT), followed by incubation in target retrieval solution for 5 minutes at 95°C and Protease III treatment (30 minutes at 40°C). Probes of interest (see Table S2) were hybridized for 2 h at 40°C and signal amplified through AMP1, AMP2 and AMP3 probes before being developed using the appropriate HRP signal. Finally, the presence of GFP was revealed using a standard immunohistochemistry protocol. Slices were blocked 30 minutes in 1% BSA – PBS 1X and incubated overnight at 4°C with the anti-GFP primary antibody (see Table S2) in 1% BSA – PBS 1X. The next day, the secondary antibody (see Table S2) was applied for 1 h at RT and nuclei stained with Hoechst (1:5000 in PBS 1X, 15 minutes at RT).

### Confocal images acquisition and analysis

An Upright Leica DM6 CS laser scanning confocal microscope was used to acquire images of the DG for each brain slice. All slides were acquired at 63X magnification with a resolution of 512 × 512, speed of 400 Hz, airy unit between 0.3 and 0.4 AU and line averaging of three. Channels were acquired from the longest wavelength to the shortest wavelength and acquired one at a time to avoid leakage between the channels with spectrum overlaps assessed with BD spectrum viewer (BD Biosciences). A tilescan of the acquired area was then stitched within the Leica LAS-X software. Images were analyzed using QuPath (v0.2.3 to v0.4.3)^28^. For each confocal image of a single DG, detected nuclei were first classified as part of the DG granular cell layer or not using a machine-learning based algorithm developed in house starting from the StarDist library. Next, nuclei were defined as positive for each given channel if their mean signal was higher than a manually pre-set threshold of detection. Finally, measurements from all detected nuclei were exported and analysed using Phyton scripts. To compute *Arc* expression levels for the triple positive nuclei (sgRNA^+^, *dCas9*^+^, *Arc*^+^), the mean intensity value corresponding to the *Arc* probe signal for each nucleus in a given DG was extracted and averaged to obtain a single value per acquired image. Overall, a total of four images was analysed for each mouse, and three to four mice per group were compared in each experiment.

### Statistics and reproducibility

Statistical analysis was performed in Prism 9 (GraphPad) and all data are displayed as mean ± standard error mean (SEM). For *in vivo* experiments, no statistical methods were used to predetermine sample sizes, but the number of animals used in each experiment is similar to those reported in previously published engram studies. Data distribution was assumed to be normal, even if this was not formally tested. Animals were randomly assigned to experimental groups and age matched. For *in vitro* experiments, at least three independent experiments were performed for statistical analysis unless otherwise specified in the figure legends. The statistical test used and the definition of n are described in the figure legends. Statistical analysis details for each figure are reported in Table S3.

**Table S1.**

Differential enrichment analysis results for the dCas9-KRAB-MeCP2 ChIP-seq experiment.

**Table S2.**

List of materials, reagents and software used in this study.

**Table S3.**

Statistics summary for all the experiments in this study.

## Acknowledgments

We thank the EPFL Bioimaging platform (BIOP) for their support in image acquisition and analysis, the EPFL Histology Core Facility (HCF) for help with the RNAScope experiments, the EPFL Gene Expression Core Facility (GECF) for performing next-generation sequencing, and the EPFL Center of Phenogenomics (CPG) for ensuring the animal welfare. We are grateful to the members of the Gräff lab for comments on the manuscript.

## Author contributions

D.M.C. and J.G. designed and conceptualized this study. D.M.C. carried out the experiments and analyzed the data. L.W. and C.L.S contributed to viral injections, behavioral experiments, histology and image analysis. L.W., M.B. and C.L.S. contributed to *in vitro* experiments. L.G. contributed to mouse colony maintenance and lentivirus production. A.M.B. performed ChIP-seq data analysis. D.M.C. and J.G. wrote the manuscript with comments from all authors.

## Data and materials availability

The list of materials, reagents and software used in this study is provided in Table S2. ChIP-seq data have been deposited in the Gene Expression Omnibus (GEO) database, https://www.ncbi.nlm.nih.gov/geo (accession no. GSE256419), all other data are available in the main text or in the supplementary materials.

---

**Extended Data Figure 1.**
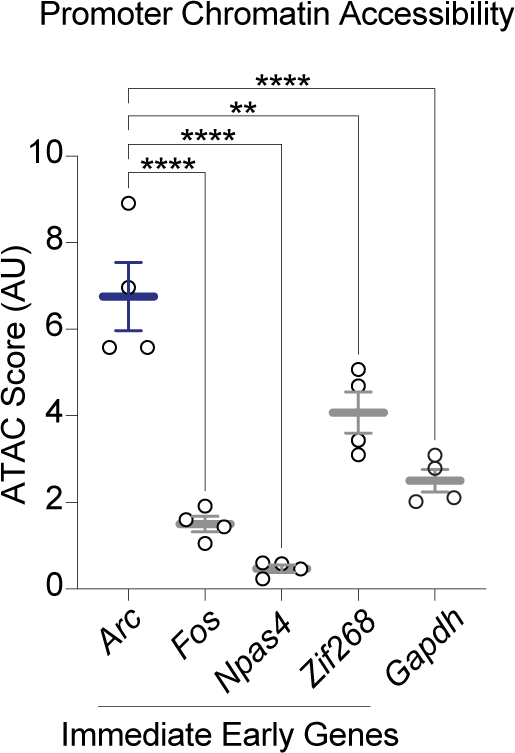
*Arc* promoter in DG cells shows increased chromatin accessibility. For DG cells from home cage mice, plotted are the RPKM values for the ATAC-seq peaks detected at the promoter regions of representative immediate early genes (*Arc, Fos, Npas4, Zif268*) and housekeeping gene (*Gapdh*). Note that the *Arc* promoter displays an highly open chromatin conformation in the baseline state compared to other immediate early genes. DG, Dentate Gyrus. Data are means ± SEM of three independent experiments compared by one-way ANOVA with Tukey’s post hoc test (**P < 0.01 and ****P < 0.0001). Data were obtained from Marco et al.^7^, GSE152954_ATAC_Desq2_Matrix_all_groups_RPKM.txt.gz.

**Extended Data Figure 2.**
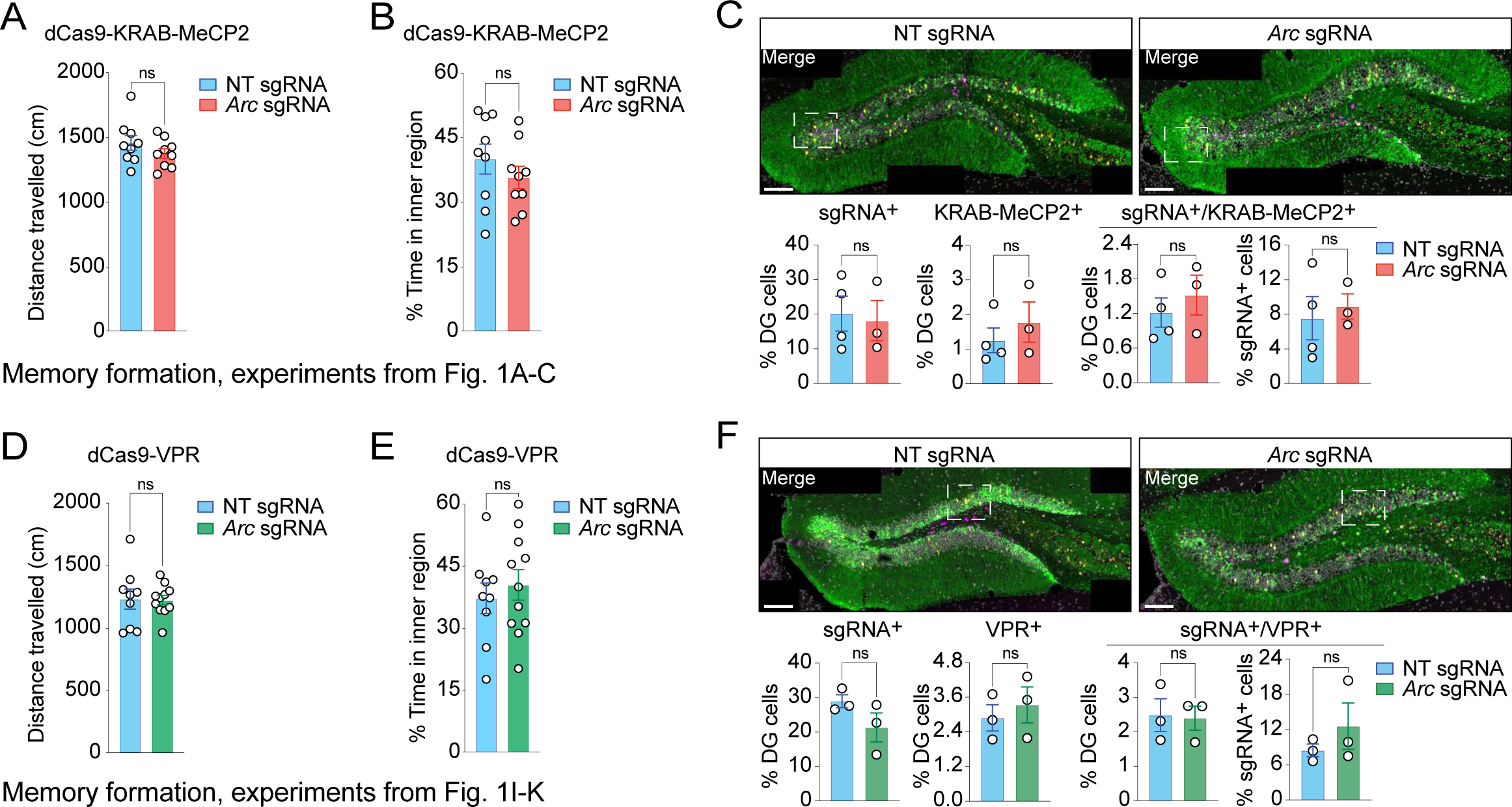
*In vivo* CRISPR-dCas9-based epigenetic editing of the *Arc* promoter does not impact basal locomotion and anxiety levels. (**A-B**) For mice from the dCas9-KRAB-MeCP2 experiment of Fig. 1A-C, displayed are (A) the average distance travelled and (B) the time spent in the inner region of the conditioning chamber during the three minutes habituation phase of the behavioral conditioning. (**C**) Top pannel: for the dCas9-KRAB-MeCP2 experiment of Fig. 1A-C, confocal images showing expression levels of *Arc* mRNA (yellow) and dCas9-KRAB-MeCP2 mRNA (magenta) alongside with NT or *Arc* sgRNA (GFP, green) in the DG. Dotted squares delimitate the areas corresponding to the enlarged images in Fig.1D. Scale bars, 100 μm. Bottom pannel: quantification of C. (**D-E**) As in A-B, but for the dCas9-VPR experiment of Fig. 1I-K. (**F**) As in C, but for the dCas9-VPR experiment of Fig. 1I-K. All data were compared by a two-tailed, unpaired t tests (ns, not significant) and plotted as means ± SEM.

**Extended Data Figure 3.**
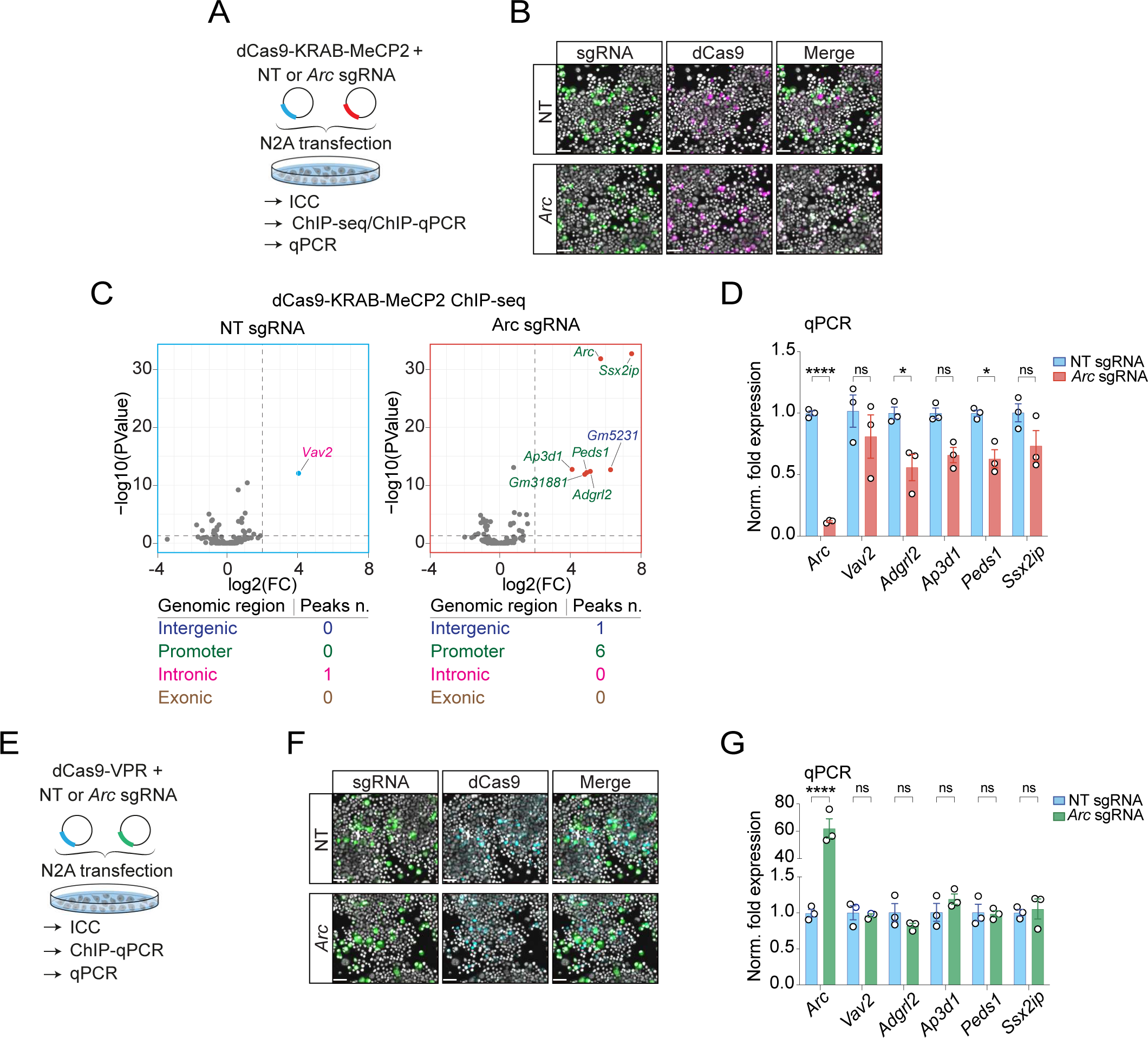
CRISPR-dCas9-based epigenetic editing enables the bidirectional control of *Arc* promoter activity with few off-target effects. (A) N2A cells were co-transfected with constructs for dCas9-KRAB-MeCP2 and *Arc* or NT sgRNA, and after 48 h collected for immunocytochemistry (ICC), ChIP-seq/ChIP-qPCR and qPCR analyses. (**B**) Representative ICC images of N2A cells treated as in A, revealing high co-transduction efficiency of sgRNA (co-expressing GFP) and dCas9-KRAB-MeCP2 (FLAG-tagged). Cell nuclei are stained with DAPI. Scale bars, 50 um. (**C**) For the samples described in Fig.1F, volcano plots showing genomic sites significantly enriched for dCas9-KRAB-MeCP2 chromatin binding (p-value < 0.05, fold change > 2). Tables below the graphs summarize the peaks’ features for each sample and dots in D are color-coded accordingly. One and 6 off-targets were identified for the NT sgRNA and *Arc* sgRNA, respectively. Of these sites, two of them were associated to predicted genes (*Gm5231* and *Gm31881*) whilst the other 5 were assigned to genes that either show low expression levels (according to the Allen Mouse Brain Atlas) in adult DG (*Vav2*, *Adgrl2* and *Peds1*), or have not been implicated in fear memory processes (*Ap3d1*, *Ssx2ip*). The differential enrichment analysis results are reported in Table S1. (**D**) For the experiment described in A, q-PCR analysis was performed for the protein coding genes identified in C. dCas9-KRAB-MeCP2 plus *Arc* sgRNA downregulated *Arc* mRNA levels compared to the control condition with minimal off-target effects. Data are means ± SEM of three independent experiments compared by two-way ANOVA with Sidak’s multiple comparisons test. (**E**) As in A, except that dCas9-VPR is expressed instead. (**F**) Representative ICC images of N2A cells treated as described in E, showing high co-transduction efficiency of sgRNA (co-expressing GFP) and dCas9-VPR (FLAG-tagged). Cell nuclei are stained with DAPI. Scale bars, 50 um. (**G**) For the experiment described in E, q-PCR analysis was performed for the protein coding genes identified in C. dCas9-VPR plus *Arc* sgRNA upregulated *Arc* mRNA levels compared to the control condition with no off-target effects. Data are means ± SEM of three independent experiments compared by two-way ANOVA with Sidak’s multiple comparisons test. For all figure panels: ns, not significant; *P < 0.05 and ****P < 0.0001.

**Extended Data Figure 4.**
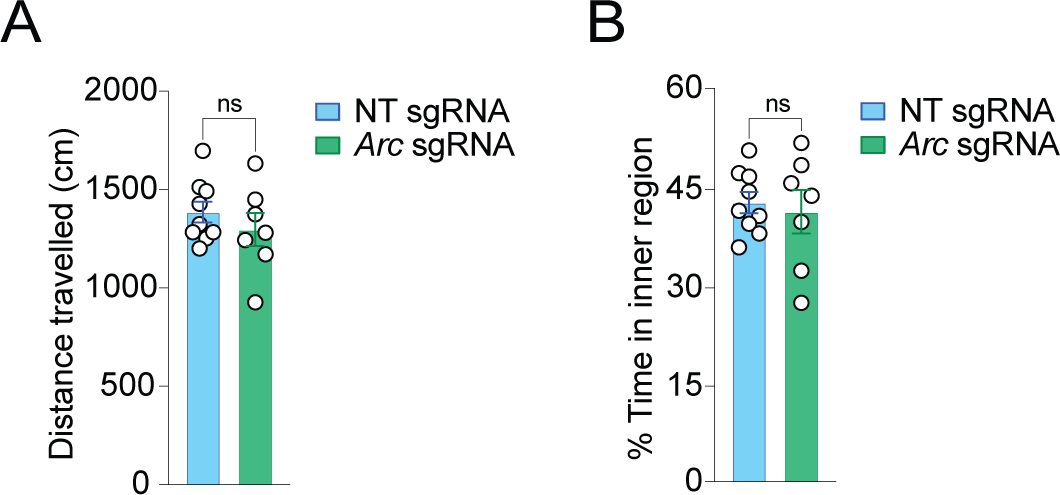
Context only control experiment: dCas9-based epigenetic editing of the *Arc* promoter does not impact basal locomotion and anxiety levels. (**A-B**) For mice from the dCas9-VPR experiment of Fig. 2H-I, displayed are (A) the average distance travelled and (B) the time spent in the inner region of the conditioning chamber during the three minutes habituation phase of the behavioral conditioning. Data were compared by a two-tailed, unpaired t test (ns, not significant) and plotted as means ± SEM.

**Extended Data Figure 5.**
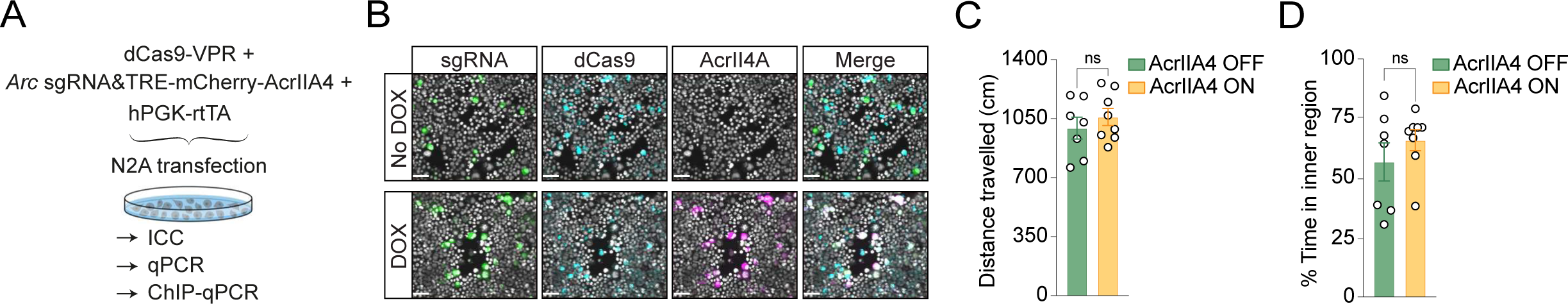
AcrllA4-dependent rescue of dCas9-VPR mediated memory enhancement does not impact basal locomotion and anxiety levels. **(A)** Experimental design to test the effect of AcrllA4 *in vitro* in the presence of dCas9-VPR and *Arc* sgRNA. 48 h after transfection, N2A cells were collected for immunocytochemistry (lCC), qPCR (Fig. 3C) and ChlP-qPCR (Fig. 3D). (**B**) Representative images of N2A cells treated as described in A, showing DOX-inducible AcrllA4 expression (mCherry reporter) and high co-transduction efficiency of *Arc* sgRNA (co-expressing GFP) and dCas9-VPR (FLAG-tagged). Cell nuclei are stained with DAPl. Scale bars, 50 um. **(C-D)** For mice from the AcrllA4 experiment of Fig. 3H-l, displayed is (C) the average distance travelled and (D) the time spent in the inner region of the conditioning chamber during the three minutes habituation phase of the behavioral conditioning. ln C and D data were compared by a two-tailed, unpaired t test and plotted as means ± SEM (ns, not significant). DOX, doxycycline.

**Extended Data Figure 6.**
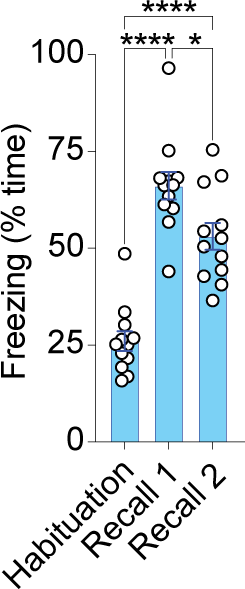
Memory retention naturally decreseases over time. Freezing levels for cFos-tTA mice (n = 12) that underwent the same CFC protocol described in Figure 2 followed by two recall sessions, the first one 72 h after CFC and the second one on the following day. Note that animals spent significantly less time freezing at the Recall 2 timepoint compared to the Recall 1 timepoint. Data are means ± SEM compared by one-way ANOVA with Tukey’s post hoc test (*P < 0.05 and ****P < 0.0001).

## References

1. Josselyn, S. A., Köhler, S. & Frankland, P. W. Finding the engram. Nat. Rev. Neurosci. 16, 521–534 (2015).

2. Choi, J.-H. et al. Interregional synaptic maps among engram cells underlie memory formation. Science 360, 430–435 (2018).

3. Liu, X. et al. Optogenetic stimulation of a hippocampal engram activates fear memory recall. Nature 484, 381–385 (2012).

4. DeNardo, L. A. et al. Temporal evolution of cortical ensembles promoting remote memory retrieval. Nat. Neurosci. 22, 460–469 (2019).

5. Coda, D. M. & Gräff, J. From cellular to fear memory: An epigenetic toolbox to remember. Curr. Opin. Neurobiol. 84, 102829 (2024).

6. Halder, R. et al. DNA methylation changes in plasticity genes accompany the formation and maintenance of memory. Nat. Neurosci. 19, 102–110 (2016).

7. Marco, A. et al. Mapping the epigenomic and transcriptomic interplay during memory formation and recall in the hippocampal engram ensemble. Nat. Neurosci. 23, 1606–1617 (2020).

8. Josipović, G. et al. Antagonistic and synergistic epigenetic modulation using orthologous CRISPR/dCas9-based modular system. Nucleic Acids Res. 47, 9637–9657 (2019).

9. Yeo, N. C. et al. An enhanced CRISPR repressor for targeted mammalian gene regulation. Nat. Methods 15, 611–616 (2018).

10. Dominguez, A. A. et al. CRISPR-Mediated Synergistic Epigenetic and Transcriptional Control. CRISPR J. 5, 264–275 (2022).

11. Reijmers, L. G., Perkins, B. L., Matsuo, N. & Mayford, M. Localization of a stable neural correlate of associative memory. Science 317, 1230–1233 (2007).

12. Shepherd, J. D. & Bear, M. F. New views of Arc, a master regulator of synaptic plasticity. Nat. Neurosci. 14, 279–284 (2011).

13. Plath, N. et al. Arc/Arg3.1 is essential for the consolidation of synaptic plasticity and memories. Neuron 52, 437–444 (2006).

14. Klemm, S. L., Shipony, Z. & Greenleaf, W. J. Chromatin accessibility and the regulatory epigenome. Nat. Rev. Genet. 20, 207–220 (2019).

15. Burns, A. M. et al. The HDAC inhibitor CI-994 acts as a molecular memory aid by facilitating synaptic and intracellular communication after learning. Proc. Natl. Acad. Sci. 119, e2116797119 (2022).

16. Marino, N. D., Pinilla-Redondo, R., Csörgő, B. & Bondy-Denomy, J. Anti-CRISPR protein applications: natural brakes for CRISPR-Cas technologies. Nat. Methods 17, 471–479 (2020).

17. Dow, L. E. et al. Conditional Reverse Tet-Transactivator Mouse Strains for the Efficient Induction of TRE-Regulated Transgenes in Mice. PLoS ONE 9, e95236 (2014).

18. Robison, A. J. & Nestler, E. J. Transcriptional and epigenetic mechanisms of addiction. Nat. Rev. Neurosci. 12, 623–637 (2011).

19. Weaver, I. C. G. et al. Epigenetic programming by maternal behavior. Nat. Neurosci. 7, 847–854 (2004).

20. Cavalli, G. & Heard, E. Advances in epigenetics link genetics to the environment and disease. Nature 571, 489–499 (2019).

21. Naik, S. & Fuchs, E. Inflammatory memory and tissue adaptation in sickness and in health. Nature 607, 249–255 (2022).

22. Netea, M. G. et al. Trained immunity: A program of innate immune memory in health and disease. Science 352, aaf1098 (2016).

23. Concordet, J.-P. & Haeussler, M. CRISPOR: intuitive guide selection for CRISPR/Cas9 genome editing experiments and screens. Nucleic Acids Res. 46, W242–W245 (2018).

24. Duke, C. G., et al. An Improved CRISPR/dCas9 Interference Tool for Neuronal Gene Suppression. Front. Genome Ed. 2, 9 (2020).

25. Sanchez-Mut, J. V. et al. PM20D1 is a quantitative trait locus associated with Alzheimer9s disease. Nat. Med. 24, 598–603 (2018).

26. Coda, D. M. et al. Distinct modes of SMAD2 chromatin binding and remodeling shape the transcriptional response to NODAL/Activin signaling. eLife 6, e22474 (2017).

27. Dixsaut, L. & Gräff, J. Brain-wide screen of prelimbic cortex inputs reveals a functional shift during early fear memory consolidation. eLife 11, e78542 (2022).

28. Bankhead, P. et al. QuPath: Open source software for digital pathology image analysis. Sci. Rep. 7, 16878 (2017).

